# Diel Oxygen Changes in Closed Ecological Systems; Predictive of Lake Metabolism?

**DOI:** 10.1101/2020.07.16.207522

**Authors:** Frieda B. Taub, David J. Bridges

## Abstract

The net oxygen change over a 24-hour day/night cycle in a laboratory study showed strong consistent patterns of (1) **gain**, when nutrients and light were available; (2) **maintain**, with daytime gains being matched by nighttime losses; and (3) **loss**, over brief periods of time during intense zooplankton grazing on previously grown phytoplankton or over long durations without an external source of nutrients. These were simplified aquatic communities closed to the atmosphere, Closed Ecological Systems (CES). Natural lakes are much more complex. While temperate lakes, having a winter accumulation of nutrients followed by sequential algal and zooplankton blooms, may show similar patterns, tropical and flood lakes may exhibit different patterns. Examination of archived lake metabolic studies could yield new insights while looking for these patterns by examining net ecosystem production (NEP), often measured as changes in oxygen concentrations.

## Introduction

It has long been known that O_2_ changes within organisms and ecosystems signify energy changes, but discerning the underlying controlling factors in complex systems has proven difficult. Patterns seen in simplified laboratory ecosystems suggest hypotheses that could be tested on archived lake data. Such an effort could yield new insights on how lake ecosystems work and gain greater understanding from the efforts used to collect, store, and analyze the information. This paper presents some patterns that have been consistently observed in Closed Ecological Systems (CES) composed of algae, with and without grazers *(Daphnia),* and with associated microorganisms. If some lakes, but not others, show similar patterns, it would increase our understanding of lake bioenergetics.

Changes in O_2_ and/or CO_2_ have long been used to estimate metabolic activity in organisms (Kleiber 1961). Odum (1956) extended O_2_ changes to aquatic community metabolism, and most of his concepts are still being used (Hoellein et al. 2013; Staehr et al. 2010). Oxygen increases are used as a surrogate for the production of organic matter, and oxygen loss as a surrogate for organic loss, after accounting for physical processes such as atmospheric exchange and possible mixing. On a daily basis, aquatic ecosystems gain, maintain, or lose organic energy, represented by net ecosystem production (NEP), usually estimated by diel O_2_ changes. Organic energy can be gained by photosynthesis, organic inputs, and chemosynthesis. Alternatively, organic energy can be lost by respiration, anaerobic metabolism, and burial. An important finding of lake metabolism is that many lakes are heterotrophic, not only being supported by in situ photosynthesis but also by terrestrial organic inputs (Cole 2013; Cole et al. 2000; Wilkinson et al. 2013).

Lake metabolism has been the focus of many current studies, but the processes driving O_2_ changes in lakes have proven difficult to identify (Coloso et al. 2008; Coloso et al. 2011a; Coloso et al. 2011b; Dugan et al. 2016; Rose et al. 2014). These processes include both biological factors, such as organism migrations and emergence or epilimnetic oxidations of hypolimnetic products of anaerobic metabolism (e.g., methane), and physical factors, such as atmospheric exchanges, precipitation, watershed drainage, longitudinal flow, vertical mixing, subsurface exchanges, and sediment burial. In addition to these processes, the properties of the mathematical models used to process the data can alter the conclusions (Dugan et al. 2016).

Numerous studies of lake O_2_ measurements have been archived in publicly available databases, including the Global Lake Ecological Observatory Network (GLEON) (Hanson et al. 2016) and Networking Lake Observatories in Europe (NETLAKE). Had the 2020 ASLO meeting not been cancelled because of the COVID-19 pandemic, there would have been two symposia set up to evaluate archived databases such as EPA National Aquatic Resource Surveys (NARS), the USGS National Water Quality Assessment (NAWQA), the National Ecological Observatory Network (NEON), the LAke multi-scaled GeOSpatial and temporal database (LAGOS), and the US Long Term Ecological Research Network (US LTER). Although available, big data are not trivial to use and are best studied by scientists familiar with the environment and its stressors.

Using the behavior of simple systems to unravel the behavior of more complex systems is a common approach. Closed Ecological Systems provide a more direct method to estimate O_2_ and CO_2_ by eliminating the need to estimate exchanges with the open atmosphere. Most aquatic studies ignore gaseous exchanges, although CO_2_ and O_2_ are among the major chemical reactants. Early CES research focused on aquatic ecology or Biological Life Support Systems (Nelson et al. 1993; Taub 1974). Rillig and Antonovics (2019) have suggested that Closed Ecological Systems are particularly useful in understanding microbial interactions.

Laboratory microcosms lack many of the components of natural lakes (Carpenter 1996; Schindler 1998), but given the rich resources of archived lake data and its complexity of interpretation, it may be fruitful to examine whether phenomena seen in these CES studies occur in some lakes and not others. Earlier efforts to generalize lake ecology found common patterns in a Plankton Ecology Group (PEG) model that suggested many temperate lakes had an annual sequence of nutrient accumulation during winter, followed by a spring algal bloom, eliminated by a zooplankton bloom, which in turn was diminished by predation; controlling factors changed as the season progressed (Lampert and Sommer 2007; Sommer et al. 1986). The experiments described here conformed to the algal-grazer pattern. Tropical lakes may not experience an annual accumulation of nutrients with a predicted sequence, and flood lakes may have dynamics driven by wet/dry cycles; these might not show similar patterns.

The purpose of this study was to explore the O_2_ dynamics among a community of algae, grazers, and microbes under various nutrient regimens and isolated from atmospheric exchange. Given a variety of initial nutrient conditions (e.g., C:N ratios), strong, constant patterns emerged that, at least superficially, seem similar to lake patterns. The consistency of the patterns suggested comparisons with archived lake data, though not an easy task, could increase our understanding of lake metabolism.

## Methods

### Organisms

Three species of green algae, *Ankistrodesmus* sp., *Scenedesmus obliquus,* and *Selenastrum capricornutum,* were grown separately in T82 medium (Astm 2012)(see also Table S1 in Supplement). *Daphnia magna* were grown on these algae in a chemically defined medium, B/4 (Taub and Mclaskey 2014); see also Table S1. The algal cultures were unialgal, but they and the *Daphnia* cultures were not free of smaller microorganisms. Therefore, all respiration measurements included microbial processes as well as algae and grazers.

### Medium

Four experiments were conducted with different nominal concentrations (in mM) of C:N:P. In Experiments 1 and 2, C 3.3, N 0.125, P 0.01, and in Experiments 3 and 4, C 13.2, N 0.031, P 0.0025 initially, with N and P increased to N 0.5, P 0.04 on day 41 (Experiment 3) and day 48 (Experiment 4). The base medium was the same in all experiments, having the concentrations of B/4 (Taub and Mclaskey 2013); see also Table S1. In general, concentrations of ions are varied by factors of 4. The concentrations were slightly altered when 100 ml of mixed algal suspension in T82 medium was added (see Tables S1 and S2). These media approximate a modified Redfield Ratio (C 106: N 16: P 1) or normalized to nitrogen (C 6.625: N 1: P 0.0625), with the quantity of carbon being doubled because only approximately half of the carbon in NaHCO_3_ would be available for photosynthesis, the rest becoming CO_3_^-2^ (Stumm and Morgan 1996), which is not available to green algae (Falkowski and Raven 2007). Phosphate was increased slightly to prevent it from being the limiting nutrient, and N:P ratios were kept constant.

### Monitoring

Temperature, pressure, O_2_, pH, and conductivity were recorded each 5 minutes by a sonde (Manta2, Eureka Water Probes) inserted into a sealed culture container; the unit was mounted on a rotary shaker at 100-110 RPM. Total O_2_ concentrations for the liquid and gas phases were calculated (Taub and Mclaskey 2013). The in vivo fluorescence of the settled algae was measured on a plant stress meter (PSM Mark II, BioMonitor SCI AB, Sweden). The *Daphnia* were counted manually. More monitoring detail is provided in the Supplement.

### Experimental Setups

A medium of the nominal concentration was prepared, and 10% of an algal mixture grown in T82 was added (e.g., 1 liter of nominal medium + 100 ml of algal mixture in T82; modified chemical compositions are calculated in Table S1). A mixture of the medium plus mixed algal suspension (850 ml) was placed in the sonde cylinder (leaving a measured gas volume). To Experiments 2 and 4, six *Daphnia* were added. The systems were sealed from the atmosphere and then placed on a platform rotating at 100 RPM.

### Physical conditions

Temperatures were nominally 20° C, and light intensities were 3.0 W/m^2^ for Experiments 1 and 2 and 6.3 W/m^2^ for Experiments 3 and 4.

## Results

### Experiment 1

In an ungrazed CES of algae and associated microorganisms, algae increased in abundance to a limit (Fig. 1A). Initially, the total O_2_ increased, as each daytime O_2_ production was slightly greater than following nighttime losses, but eventually diel O_2_ changes became smaller and in closer agreement (Fig. 1B).

**Fig. 1.**
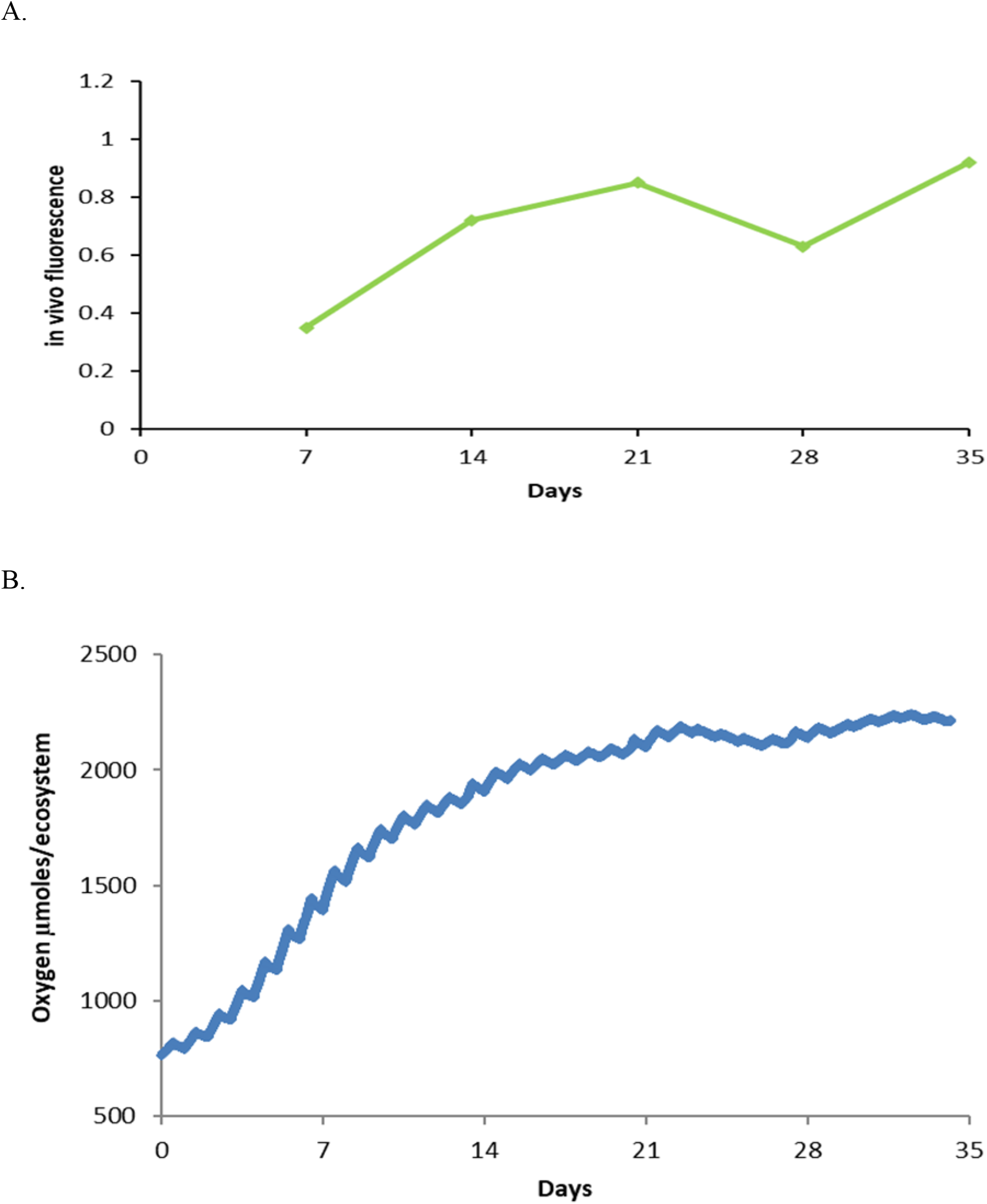
An ungrazed Closed Ecological System consisting of mixed algae and associated microorganisms. **A.** increase in algal density (in vivo fluorescence); the initial value was too low to measure on the PSM instrument. **B.** Oxygen changes during 12:12 hour light/dark cycles with the exception of days 23-25 (8:16) and days 29-31 (18:6). Initially, O_2_ increased to a greater degree than it decreased in the dark with a net diel gain; later the increases and decreases were in close agreement, a period of maintenance. Note that all of the B figures have the same vertical axis of 500-2500 μmoles per ecosystem so they can be compared.

### Experiment 2

In a grazed CES of mixed algae, *Daphnia,* and associated microorganisms, algae increased initially, but were eventually reduced in abundance as the *Daphnia* population developed, followed by “predator-prey dynamics” between the algae and *Daphnia* (Fig. 2A). Initially, the total O_2_ increased sharply, as daytime O_2_ production was greater than the following nighttime losses. Eventually, however, as the *Daphnia* population increased to a maximum on day 14, the O_2_ gain was terminated, followed by a loss, with the O_2_ increase being less than the following night’s decrease (Fig. 2B). When the *Daphnia* had reduced the algal abundance, the *Daphnia* population decreased and allowed a brief period of gain and eventual maintenance.

**Fig. 2.**
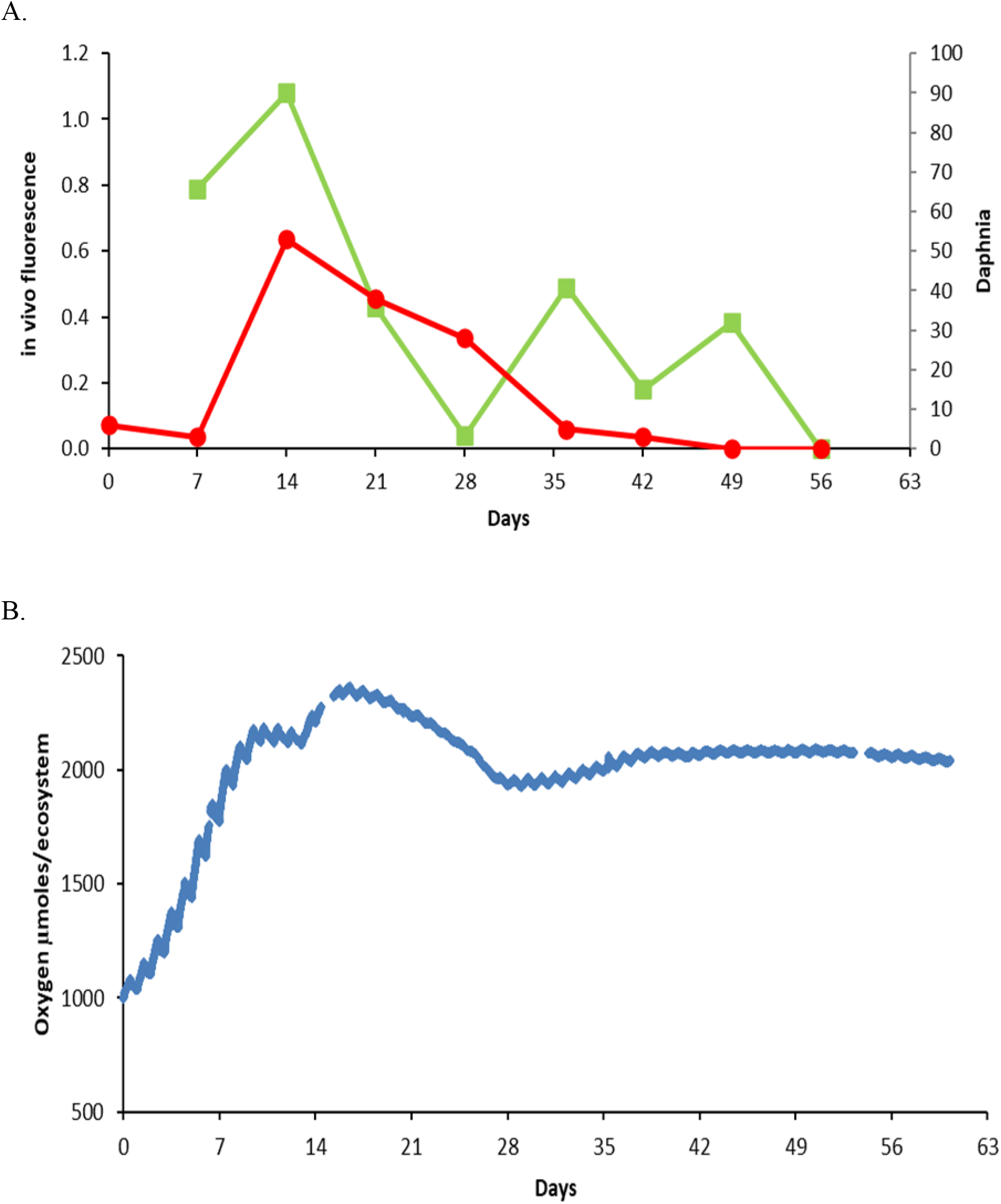
A grazed Closed Ecological System consisting of mixed algae, *Daphnia,* and associated microorganisms. **A.** Algal abundance (in vivo fluorescence). The value at time 0 was too low to measure. **B.** Oxygen dynamics. Note the earlier period of gain, while daytime O_2_ production was greater than the following nighttime’s loss (see changes in light/dark cycle days 10-15)* until the *Daphnia* grazing reduced the algal abundance, associated with decreases in total O_2_, the gain during the daytime being less than the following nighttime’s loss. Eventually, the ecosystem maintained a fairly constant total O_2_ as the small daytime gains and nighttime losses were similar. *Short days 10-12, long days 13-15 (data lost for days 14 and 15). Medium was C 3.3, N 0.125, P 0.01 mM. [2011Sept28_NaHCO3_Dap_M2 Charts *Daphnia* &; PSM chart, Total O_2_]

### Experiment 3

In an ungrazed CES of algae and associated microorganisms, to which limiting nutrients were later added, algal abundance increased to a limit and then started to decline; subsequently, N and P were added, and abundance increased slightly (Fig. 3A). The total O_2_ (Fig. 3B) increased during a period of gain, production being slightly greater than the following nighttime loss, but eventually the total O_2_ stabilized as gains and losses were similar and then decreased as the daytime gain was slightly less than the nighttime loss. Nutrients were added on day 41 (requiring the CES to be opened, resulting in an O_2_ drop), after which there was a marked increase in total O_2_ as gains exceeded losses.

**Fig. 3.**
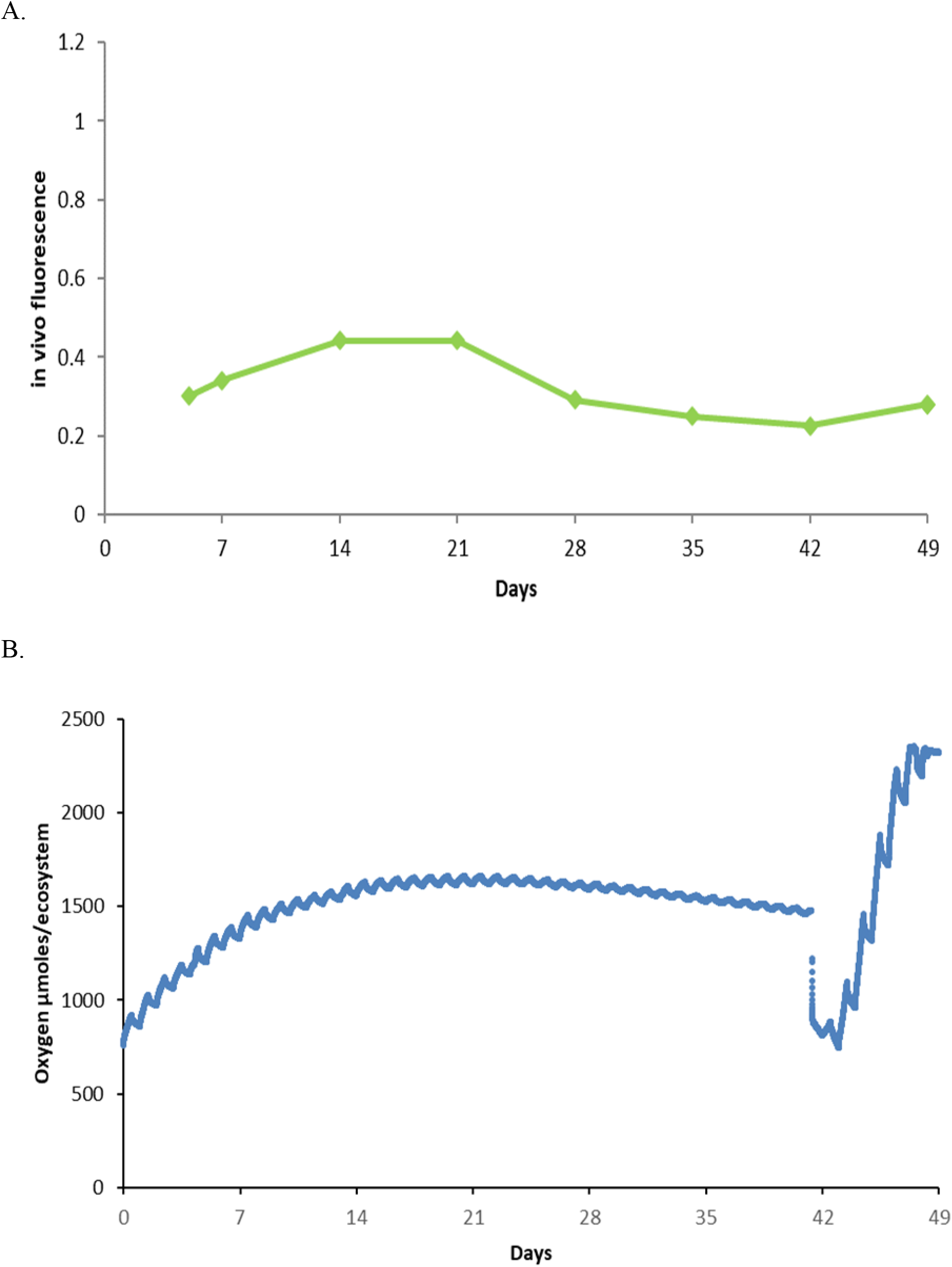
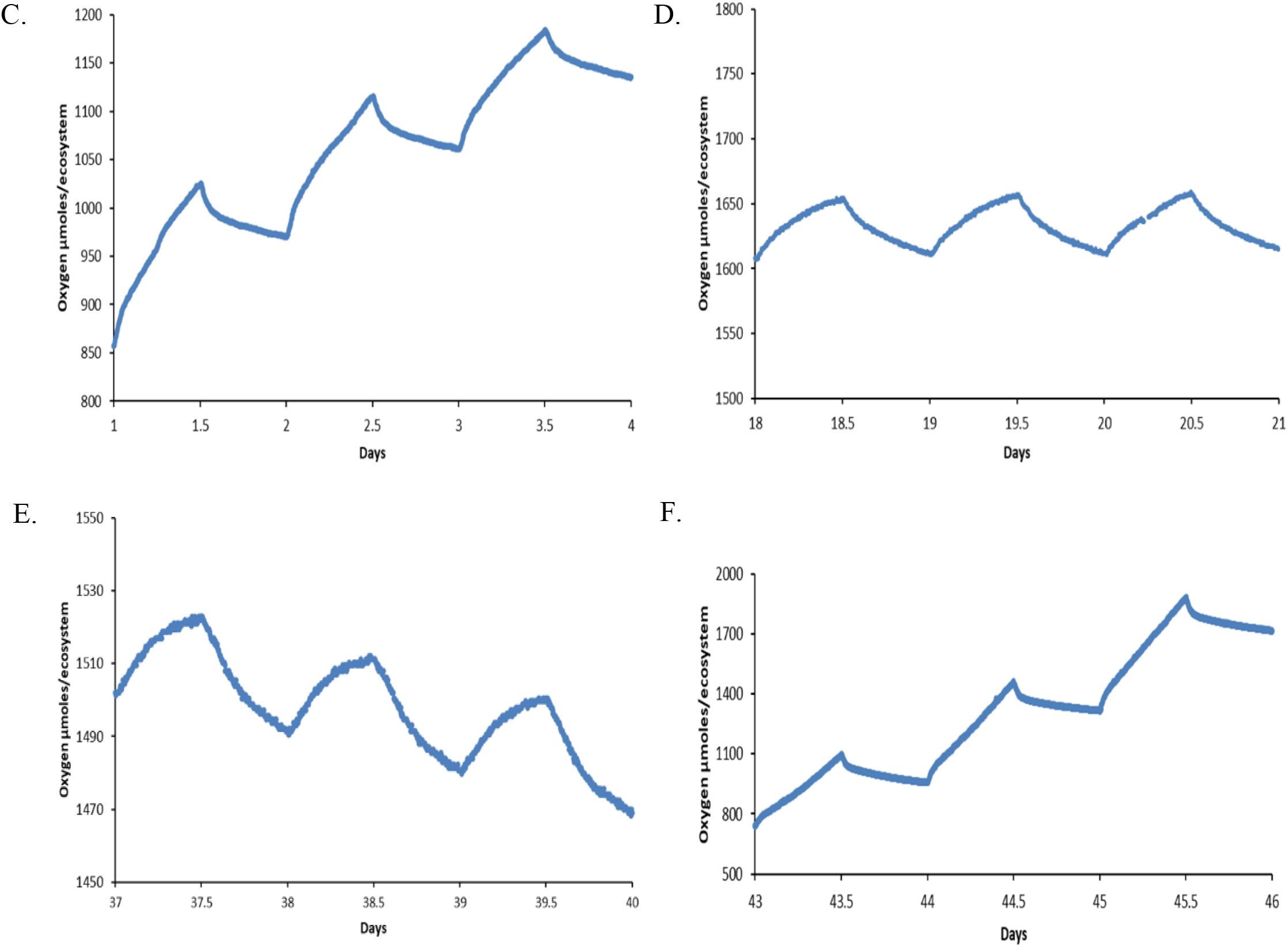
An ungrazed Closed Ecological System consisting of algae and associated microorganisms with N and P addition on day 41. **A.** Algal abundance (in vivo fluorescence); the value at time 0 is too low to measure. **B.** Oxygen dynamics showing periods of O_2_ gain, maintenance, loss, and (after nutrient enrichment) gain. **C.** Details of O_2_ change: days 1-4 (Gain); **D.** days 18-21 (Maintain); **E.** days 37-40 (Lose); **F.** after enrichment, days 43-46 (Regain). For the 3-day charts (**C-F**), note that the range of the y axis changes. Initially C 13.2, N 0.03125, P 0.0025; after enrichment on day 41, N 0.5, P 0.01 mM (Nominal).

The detailed shapes of the O_2_ dynamics during the light and dark phases were nonlinear. The increase in O_2_ was fastest during the first hour of light, and the decrease was fastest during the first hour of dark (Fig. 3C, D, E). During the initial period of diel gain, both increases and decreases were large, increases being only slightly greater than decreases (Fig. 3C). During maintenance, the changes were smaller but equal (Fig. 3D). During losses, changes were similar with gains being only slightly less than losses (Fig. 4E). With nutrient addition, gains again exceeded losses.

**Fig. 4.**
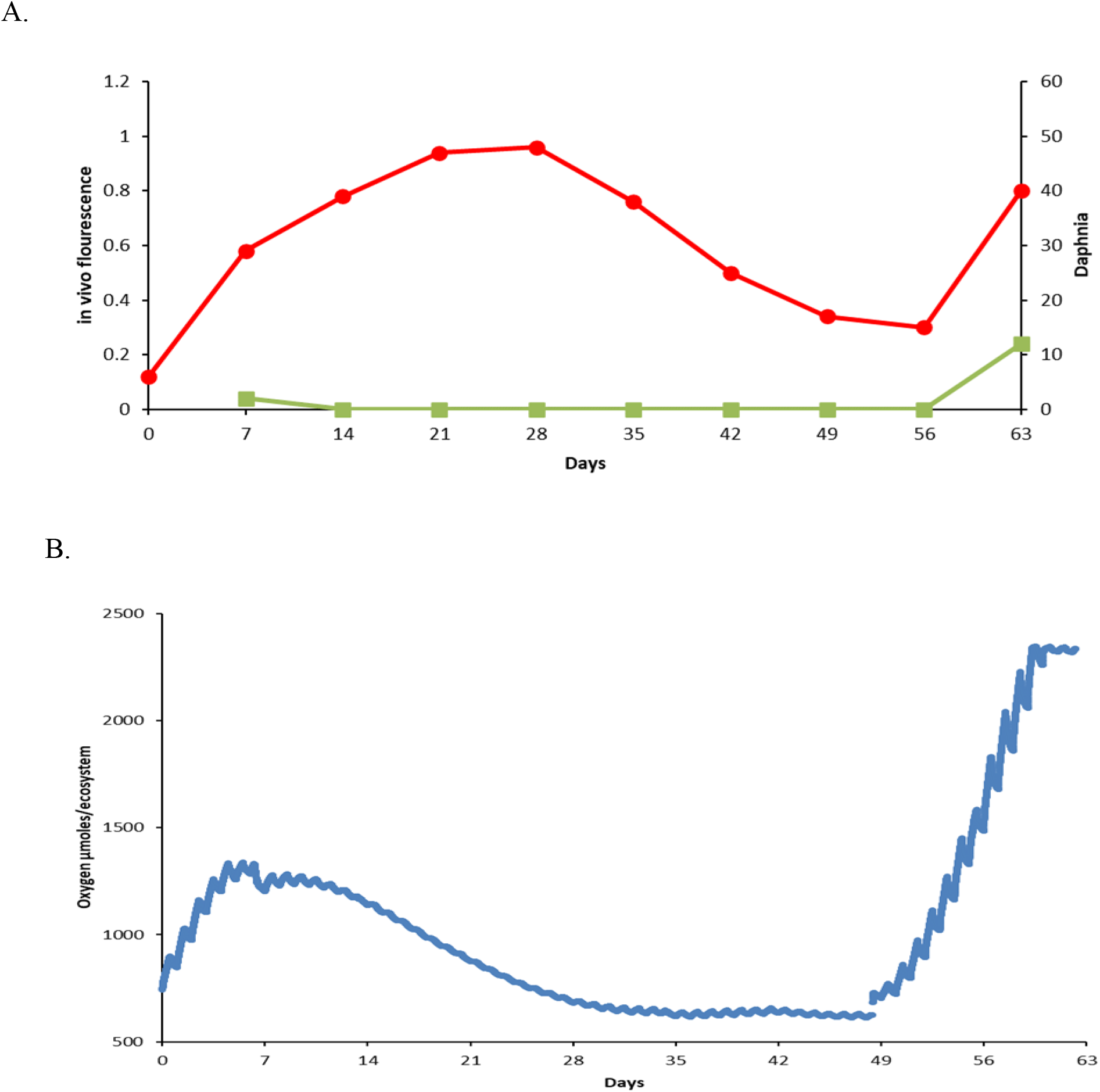
A grazed Closed Ecological System consisting of algae, *Daphnia,* and associated microorganisms with late nutrient addition on day 48. A. Algal abundance (in vivo fluorescence) (Green) was too low to measure initially and again later in the experiment until nutrient addition; *Daphnia* (Red) increased in abundance to day 28, and thereafter decreased until a week after nutrient addition. B. Oxygen dynamics initially increased, decreased as the *Daphnia* population increased and algal abundance became unmeasurable, the system was at steady state (maintenance) and after N and P addition, a period of rapid gain. Initial C 13.2, N 0.03125, P 0.0025; after addition on day 48, N 0.5, P, 0.04 mM (Nominal). [2015 Jan8_hiNaHCO_3__Daph, charts DaphGraph (3^rd^ down) and TotalO_2_]

### Experiment 4

In a grazed CES with mixed algae, *Daphnia,* and associated microorganisms, with late nutrient enrichment, the algal population was reduced to unmeasurable abundance almost immediately by the increasing *Daphnia* abundance up to day 28 (Fig. 4A), after which *Daphnia* decreased only slightly. After nutrient enrichment, the algae (in vivo fluorescence) were again measurable (Fig 4A). For oxygen dynamics, there was a period of gain for 6 days, followed by losses and eventual steady state; after N and P additions, there was a period of O_2_ gain.

## Discussion

Three findings stand out. The most surprising finding was that a period of intense grazing was associated with a diel O_2_ loss (negative NEP), which, if considered on a short-term basis, might be interpreted as heterotrophy, but was actually caused by the grazing of algae that had been produced earlier. Since these systems are closed to material exchanges, no organics could have entered. The reduced diel O_2_ production was the result of reduced algal populations. Given a reproducing *Daphnia* population, the algal density was often below our detection limits in many of our experiments. Our second finding of note is that small densities of algae, given available nutrients, could produce and use large amounts of O_2_, whereas denser, nutrient-depleted algae produced and used lessor amounts of O_2_. This could be a concern when community metabolism is estimated by multiplying organism density by a constant estimate of metabolism. Finally, there was strong linkage between O_2_ produced and O_2_ used; photosynthetically active communities had high respiration, whereas, once nutrients were sequestered, communities had both low photosynthetic and low respiratory rates.

A potential mechanism for the linkage between daytime and nighttime O_2_ changes could be that well-nourished fast-growing organisms have high metabolic rates and starved organisms often have low metabolic rates. For example, P-limited *Selenastrum,* one of our green algae, had a fivefold lower rate of respiration and a threefold reduction in CO_2_ fixation and O_2_ production, as compared with better nourished cells (Theodorou et al. 1991), and *Daphnia* that had no or limited food for 15-20 hours increased their respiratory rates three-to fourfold when fed (Schmoker and Hernandez-Leon 2003). There may be additional mechanisms that contribute to steady-state maintenance conditions, O_2_ nighttime uptake being similar to daytime O_2_ production. Even when photosynthesis has been minimized during nutrient deficiencies, the nighttime respiration was generally equal to or only slightly greater than the net produced the previous day.

Most findings conformed to expectations that a small number of algae inoculated into fresh medium would undergo a potential lag, exponential increase, stability, and decline (Fogg and Thake 1987). The overgrazing of algae by *Daphnia,* and the eventual decline of the *Daphnia* and regrowth of algae, is also a familiar “predator-prey relationship” (Sommer et al. 1986). The data presented here are part of a larger series of experiments involving different C:N ratios (Taub 2019) and a variety of organic substrates (Taub 2009) with more experiments yet to be reported.

The shape of the diel O_2_ changes, with faster rates shortly after lights on and/or off, can have several interpretations. Sadro et al. (2011) and Schindler et al. (2017) have interpreted the rapid O_2_ loss at the beginning of darkness as microbial metabolism of photosynthetically produced dissolved organic carbon. Earlier studies interpreted increased respiration after photosynthesis as algal (or plant) metabolism of carbohydrates, later becoming slower as proteins are metabolized (Falkowski and Raven 2007), but it is not certain if those samples were free of microbes. The shapes of both the rapid O_2_ increase at lights on and the rapid O_2_ decrease at lights off are similar to simple mathematical models of resource-dependent reactions, CO_2_ and possibly N and P being more abundant at lights on from metabolism during the previous night, and organic substrates being more abundant at the end of the lighted period and the beginning of darkness (lights off). The shapes are also similar to a more complex model involving production and destruction (recycling) (Odum and Odum 2000).

Comparisons with photosynthesis-respiration (P-R) rates measured in natural aquatic communities almost all show a strong linear relationship with a lot of scatter (Solomon et al. 2013). Reasons for deviations from P approximating R could include an input of organic material that would increase respiration to exceed primary production or an input of inorganic nutrients that would increase photosynthesis above respiration.

Of greatest interest would be examining if intense grazing reduces diel O_2_ production and might be interpreted as heterotrophy (if short times are considered) but actually represents consumption of earlier photosynthetic products.

The CES experiments were originally designed to simulate the PEG model in which inorganic nutrients accumulate during a winter of low light intensity and temperature and spring brings an algal bloom that is terminated (in oligotrophic systems) by zooplankton grazing (Lampert and Sommer 2007; Sommer et al. 1986). Temperate lakes that conform to the PEG model might be likely to have algal-grazer dynamics similar to a Closed Ecological System. However, it is entirely possible that other factors may obscure metabolic activity, e.g., migrations, storms causing vertical mixing and terrestrial runoff, or flushing with upstream inputs and downstream drainage. Tropical lakes might not be expected to show behavior similar to that in a CES, since nutrients do not accumulate during a cold, dark period, leading to sequential blooms of phytoplankton, grazers, and predators; rather, wet and dry seasons may drive community responses. Flood lakes have episodic inputs of terrestrial materials that have major effects on community structure and metabolism (Holtgrieve et al. 2013; Holtgrieve et al. 2010).

Hypotheses based on simple systems might be used to explore data sets of naturally complex aquatic communities. In simple Closed Ecological Systems, there is a consistent linkage between daytime O_2_ increase and the following night’s O_2_ decrease, with systems showing O_2_ levels will (1) **gain,** if nutrients and light are available; (2) **maintain**, with increases and decreases balancing (steady state); and (3) **loss**, after long periods of nutrient limitation. Do periods of high grazing in lakes have negative O_2_ changes that are generally interpreted as heterotrophy but caused by the consumption of previously grown algae? Thus laboratory microcosms, with their limitations, might increase the value of the archived lake data sets.

## Acknowledgements

Anna K. McLaskey was involved with the initial development of the sonde experiments, gaseous calculations, and Experiments 1 and 2. Irina Moruga assisted in the chemical calculations, data analyses, and preparation of the manuscript. Other laboratory assistants included Christina H. Tran, Ty Mahoney, and Samuel Turner. Editing assistance was provided by Kristie Hammond of.

