## Supplemental Material for "Diel Oxygen Changes in Closed Ecological Systems; Predictive of Lake Metabolism?"

Frieda B. Taub*^[1]^ and David J. Bridges
School of Aquatic and Fishery Sciences, College of the Environment, University of Washington, Seattle, WA 98195

**Introduction**

These studies were a small component of testing various C:N relationships for their effects on pH and *Daphnia* survival in Closed Ecological Systems (Taub 2019). In most lakes, N and/or P are the limiting factors for photosynthesis; carbon is usually considered to be amply available, being supplied by respiration and atmospheric exchange (Schindler 1974). In ecosystems isolated from the atmosphere, it is necessary to supply an inorganic source of carbon to explicitly measure O_2_ and CO_2_ exchanges. The use of NaHCO_3_ is a convenient method of controlling the initial quantity of dissolved inorganic carbon. In brief, it appears that if N is high, so much C is removed from NaHCO_3_ that the pH increases beyond the tolerance of the *Daphnia*. Whereas, if N is very low, less NaHCO_3_ is converted to CO_3_^-2^, and the *Daphnia* population survives longer. The addition of high concentrations of NaHCO_3_ to provide pH buffering was not successful. All salt concentrations seemed to be within the osmotic tolerance of *Daphnia*, so the area is still under research. A fuller set of experimental results will be published soon, and data are available if requested. However, the strong, consistent results on diel O_2_ dynamics seemed relevant to current limnological studies.

**Methods**

**Experimental design:** Considering C, N, and P as the most likely factors limiting photosynthesis, various concentrations were tested, as shown in Table S1. The other nutritional requirements, presumed nonlimiting to photosynthesis, were constant, termed CORE (Table S2). To 1 liter of the specific medium, 100 ml of mixed algae culture was added. The algae had been grown in T82 medium (4x the concentration of B/4 medium and lacking trace metals and algae required by *Daphnia*), so the experimental chemical concentrations were slightly altered (Tables S1 and S2). Because the algae were grown on atmospheric CO_2_, we do not know the precise amount of carbon the algae contributed. Based on other experiments, however, we estimate the algal cultures contributed approximately 0.63 mM particulate carbon and 0.064 mM dissolved organic carbon, which is considered trivial compared to the NaHCO_3_ added. The carbon and nitrogen contributed by the *Daphnia* are calculated in Table S3 and considered trivial. Details for making the media (master solution concentrations, volumes, etc.) are given in (Astm 2012) and are available upon request from the corresponding author.

**Organisms:** The algae consisted of *Ankistrodesmus sp*., *Scenedesmus obliquus* and *Selanastrum capicornutum*; the grazer was *Daphnia magna*. The *Scenedesmus obliquus* and *Daphnia magna* were obtained from Michael Brett (Professor of Civil & Environmental Engineering, University of Washington). These organisms have been reared for more than 20 years on chemically defined media (algae, T82, and *Daphnia* B or B/4). The cultures contain associated microorganisms, but no protozoa large enough to graze on the algae.

The chemical compositions of B/4, T82, and 1 liter of B/4 amended with 100 ml of T82 (and algae) are shown in Tables S1 and S2. The T82 medium has long been used by our laboratory to grow algal species; B medium is T82 plus trace metals and vitamins required by *Daphnia magna* (Keating 1985). Through experience, a one-quarter dilution of B medium (B/4) has proven more successful in rearing *Daphnia*. In these experiments, 100 ml of mixed algae in T82 was added to 1 liter of B/4 medium; the resultant chemical compositions are shown in Table S2. Only C, N, and P are likely to be limiting.

We used a Manta2 sonde, which has been fully described elsewhere (Taub and Mclaskey 2013). It was manufactured by Eureka Water Probes ([www.waterprobes.com](http://www.waterprobes.com), phone 512 302 4333), and modified for laboratory work by converting it from battery to line power and manufacturing an airtight, clear plastic container of approximately 1 liter, of which 150 ml was occupied by the sensors. We used the sonde mainly for measurements of dissolved O_2_, pH, and temperature. In vivo chlorophyll measurements are not indicative of the algal abundance, because most of the algae settled and were not measured by the Chl a sensor at the top of the device. The settled algae at the clear bottom was measured by a plant stress meter (BioMonitor S.C.I. AB, a Swedish brand no longer available); using the Fm reading at 5 seconds, light level 4. The sonde was mounted on a rotary shaker (Innova 2300, New Brunswick Scientific) at 100-110 RPM; in the absence of mixing, the O_2_ appears to increase after lights off because most of the photosynthesis is taking place near the bottom of the container and it takes time for the O_2_ to diffuse to the O_2_ probe near the top if mixing is inadequate.

The total O_2_ for the ecosystem was calculated from the recorded O_2_ as mg l^-1^, converted to mM by dividing by 32 (the molecular mass of O_2_), multiplying by 1000 (to convert mM to µM), and adding the O_2_ in the liquid and gas phases. Based on temperature and O_2_ solubility, the gaseous O_2_ concentration was 29.9 times that of the liquid (Taub and Mclaskey 2013). For all experiments, the liquid volume was 850 ml; for Experiments 1, 3, and 4, the gas volume was 60 ml; for Experiment 2, the gas volume was 90 ml. For 1, 3, and 4, the total O2 µmol/ecosystem = (µmol O2) *(0.85+(0.06*29.9))=2.644, and for Experiment 2, the total O2 µmol/ecosystem = (µmol O2) *(0.85+(0.09*29.9))=3.541. As a result, the concentration in the liquid (µmol) was multiplied by 2.644 for Experiments 1, 3, and 4 and by 3.541 for Experiment 2.

Light intensity was measured by Solar Light Company Model PMA2100 with PAR Detector PMA2132 SN2973; data are shown in Table S4. The light intensities, ranging from 2.7-6.6 W m-2 were low relative to most incubator experiments. Usually measured as W m^-2^, the readings can be converted to µmol quanta m^-2^ s^-1^ by multiplying by 4.6. Because the top of the Manta is black, the light measurements were taken at the midpoint of the clear section from four directions and averaged. Cool white fluorescent lights were used.

**Discussion**

By using diel O_2_ change, it is unnecessary to estimate gross primary production. Given that during the day both photosynthesis and respiration are occurring, Odum (1956) made the simplifying assumption that daytime (lighted) respiration was equal to nighttime (dark) respiration, and most researchers have continued to use that assumption (Hoellein et al. 2013; Staehr et al. 2010). This was convenient because many highly productive systems have small net ecosystem production given that both photosynthesis and simultaneous respiration are high. However, attempts to estimate photosynthesis and respiration using ^18^O have questioned that assumption (Hotchkiss and Hall 2014). Our analysis does not need that assumption since the daytime respiration is energy that has already been expended and is not available to do new chemical work of growth, etc. An alternative method of displaying our data is shown for two of the experiments discussed here (Taub and Mclaskey 2014).

Table S1. Nominal concentrations of limiting nutrients in medium before and after algae addition.

| Nominal Concentrations of C:N:P used | | |  |  |  |  | Conc. after 100 ml of algal suspension in T82 | |  |  |
| --- | --- | --- | --- | --- | --- | --- | --- | --- | --- | --- |
| **Exp. 1** | 2012May17_NaCO3_M2 | |  |  |  |  |  |  |  |  |
| Nutrients | mM | Element | mg/l |  |  |  |  | mM |  |  |
| NaHCO3 | 3.300 | C | 39.60 |  |  |  | C | > 3.3 |  |  |
| NaNO3 | 0.125 | N | 1.75 |  |  |  | N | 0.159 |  |  |
| KH2PO4 | 0.010 | P | 0.31 |  |  |  | P | 0.013 |  |  |
| **Exp. 2** | 2011Sept28_NaHCO3_Daph | |  |  |  |  |  |  |  |  |
| Nutrients | mM | Element | mg/l |  |  |  |  |  |  |  |
| NaHCO3 | 3.300 | C | 39.60 |  |  |  | C | > 3.3 |  |  |
| NaNO3 | 0.125 | N | 1.75 |  |  |  | N | 0.159 |  |  |
| KH2PO4 | 0.010 | P | 0.31 |  |  |  | P | 0.013 |  |  |
| **Exp. 3** | 2014_Sept3_hiNaCO3_2 | |  |  | Late Addition |  | Initial |  |  | Late Addition |
| Nutrients | mM | Element | mg/l |  | day 41 |  |  |  |  |  |
| NaHCO3 | 13.200 | C | 158.4 |  | mM |  | C | >13.2 |  | >13.2 |
| NaNO3 | 0.031 | N | 0.438 |  | 0.5 |  | N | 0.074 |  | 0.5 |
| KH2PO4 | 0.003 | P | 0.077 |  | 0.04 |  | P | 0.006 |  | 0.04 |
| **Exp. 4** | 2015_Jan8_hiNAHCO3_Daph | |  |  | Late Addition |  |  |  |  |  |
| Nutrients | mM | Element | mg/l |  | day 48 |  |  |  |  |  |
| NaHCO3 | 13.200 | C | 158.4 |  |  |  | C | >13.2 |  | >13.2 |
| NaNO3 | 0.031 | N | 0.438 |  | 0.5 |  | N | 0.074 |  | 0.5 |
| KH2PO4 | 0.003 | P | 0.077 |  | 0.04 |  | P | 0.006 |  | 0.04 |

Table S2. CORE, nonlimiting nutrients, common to all experiments, before and after algal additions.

| CORE Medium, all experiments. | | | | |  | Estimated after algal addition | | | |
| --- | --- | --- | --- | --- | --- | --- | --- | --- | --- |
|  | formula weight | |  |  |  | in medium T82 (4x conc. Of B/4) | | | |
| **Major Salts** |  | mM |  | mg/l |  | Major Salts | mM |  | mg/l |
| MgSO4·7H2O | 246.5 | 0.0250 | Mg | 0.608 |  | MgSO4·7H2O | 0.032 | Mg | 0.773 |
| NaOH | 40 | 0.0080 | Na | 0.185 |  | NaOH | 0.010 | Na | 0.235 |
| CaCl2·2H2O | 147 | 0.2500 | Ca | 10.000 |  | CaCl2·2H2O | 0.318 | Ca | 12.727 |
| NaCl | 58.5 | 0.3750 | Na | 8.625 |  | NaCl | 0.477 | Na | 10.977 |
| Al2(SO4)3·18H2O | 666.5 | 0.0012 | Al | 0.065 |  | Al2(SO4)3·18H2O | 0.002 | Al | 0.083 |
| Na2SiO3·9H2O | 284 | 0.0200 | Na | 0.920 |  | Na2SiO3·9H2O | 0.025 | Na | 1.171 |
|  |  |  | Si | 0.560 |  |  |  | Si | 0.713 |
| **Trace Metals** |  | µM |  | mg/l |  | Trace Metals | µM |  |  |
| FeSO4·7H2O | 278 | 1.4000 | Fe | 0.0781 |  | FeSO4·7H2O | 1.782 | Fe | 0.0994 |
| EDTA | 292 | 1.7750 | EDTA | 0.5183 |  | EDTA | 2.259 | EDTA | 0.6596 |
| H3BO3 | 61.8 | 0.9375 | B | 0.0100 |  | H3BO3 | 1.193 | B | 0.0127 |
| ZnSO4·7H2O | 287.5 | 0.0313 | Zn | 0.0019 |  | ZnSO4·7H2O | 0.040 | Zn | 0.0024 |
| MnCl2·4H2O | 197.9 | 0.3125 | Mn | 0.0169 |  | MnCl2·4H2O | 0.398 | Mn | 0.0215 |
| Na2MoO4·2H2O | 242 | 0.0313 | Mo | 0.0030 |  | Na2MoO4·2H2O | 0.040 | Mo | 0.0038 |
| CuSO4·5H2O | 249.7 | 0.0063 | Cu | 0.0004 |  | CuSO4·5H2O | 0.008 | Cu | 0.0005 |
| Co(NO3)2·6H2O | 291 | 0.0031 | Co | 0.0002 |  | Co(NO3)2·6H2O | 0.004 | Co | 0.0002 |
|  |  |  |  |  |  | (the following are not in T82) | | | |
| **Trace Metals for *Daphnia*** | | µM |  | mg/l |  | Trace Metals for *Daphnia* | | | |
| LiCl | 42.39 | 1.82500 | Li | 0.01250 |  | LiCl | 1.659 | Li | 0.01136 |
| RbCl | 120.9 | 0.14500 | Rb | 0.01250 |  | RbCl | 0.132 | Rb | 0.01136 |
| SrCl2 6H20 | 266.6 | 0.14250 | Sr | 0.01250 |  | SrCl2 6H20 | 0.130 | Sr | 0.01136 |
| NaBr | 102.9 | 0.04000 | Br | 0.00325 |  | NaBr | 0.036 | Br | 0.00295 |
| KI | 166 | 0.00500 | I | 0.00063 |  | KI | 0.005 | I | 0.00057 |
| H2SeO3 0.0016 | 129 | 0.00325 | Se | 0.00025 |  | H2SeO3 | 0.003 | Se | 0.00023 |
| Na3VO4 | 183.8 | 0.00243 | V | 0.00013 |  | Na3VO4 | 0.002 | V | 0.00011 |
| **Vitamins for *Daphnia*** | | mM |  | mg/l |  | Vitamins for *Daphnia* | | | |
| B12 (cyanocobalamin) | 1355 | 9.22E-08 | B12 | 0.00013 |  | B12 | 8.4E-08 | B12 | 0.00011 |
| Biotin (d-biotin) | 244.3 | 5.12E-07 | Biotin | 0.00013 |  | Biotin | 4.7E-07 | Biotin | 0.00011 |
| Thiamine (HCl) | 337.3 | 7.41E-05 | Thiamine | 0.02500 |  | Thiamine | 6.7E-05 | Thiamine | 0.02273 |

Table S3. Potential C, N, P addition from 6 *Daphnia* (2 small, 2 medium, 2 large). Dry weights are taken from Taub and Mclaskey (2014) using the equations from Burns (1968). Schindler (1968) had slightly lower dry weights. The C:N:P proportions are from Andersen and Hessen (1991). The weight of the element is converted to molar quantities, and then to concentrations after being added to 0.850 liter of the designated medium.

| Sources |  |  |  |  |
| --- | --- | --- | --- | --- |
|  |  | 2 of each size | |  |
|  | mg/animal | mg |  |  |
| small | 0.004 | 0.008 |  |  |
| med | 0.074 | 0.148 |  |  |
| lg | 0.218 | 0.436 |  |  |
|  | Total | 0.592 | mg *Daphnia* | |
| %C | %N | %P | from Hessen Fig. 2 | |
| 0.48 | 0.09 | 0.014 |  |  |
| 0.28416 | 0.05328 | 0.008288 |  | mg of each element |
| 0.02368 | 0.003806 | 0.000268 |  | mMoles of each element |
| 0.027859 | 0.004477 | 0.000315 | mMolar | concentration added to 0.85 l |
| This is insignificant compared to the usual media components. | | | | |

Table S4. Physical conditions recorded for each experiment.

|  |  |  |  |  | Light Intensity | | |  | Temperature | |
| --- | --- | --- | --- | --- | --- | --- | --- | --- | --- | --- |
|  |  |  |  |  |  |  | µE/m2 = 4.648 uE/m2 s |  |  |  |
|  |  |  |  |  | W/m2 |  | µE/m2 s |  | Temp |  |
| Exp. 1 | 2012May17_NaCO3_M2 | | |  |  |  |  |  |  |  |
| Light info. | | not in data file | |  |  |  |  |  |  |  |
|  | using Light Intensity 2012May3 | | | | 2.7 |  | 12.5496 |  | Mean Temp | 20.8 |
|  | using side light intensity only | | |  |  |  |  |  | St DevP | 0.1 |
|  |  |  |  |  |  |  |  |  | Min | 20.7 |
|  |  |  |  |  |  |  |  |  | Max | 20.8 |
| Exp. 2 | 2011Sept28_NaHCO3_Daph | | |  |  |  |  |  |  |  |
|  | in data file | |  |  | 3.37 |  | 15.66376 |  | Mean | 20.7 |
|  | using side only. | |  |  |  |  |  |  | St DevP | 0.1 |
|  |  |  | Mean |  | 3.0 |  |  |  |  |  |
|  |  |  |  |  |  |  |  |  | Min | 20.5 |
|  |  |  |  |  |  |  |  |  | Max | 21.4 |
| Exp. 3 | 2014_Sept3_hiNaCO3_2 | | |  |  |  |  |  |  |  |
|  | in data file, Light Intensity | | |  | 6.63 |  | 30.81624 |  | Days 0-49 (row 14218) | |
|  | using side only. | |  |  |  |  |  |  | Mean | 20.4 |
|  |  |  |  |  |  |  |  |  | StDevP | 0.2 |
|  |  |  |  |  |  |  |  |  | Min | 19.9 |
|  |  |  |  |  |  |  |  |  | Max | 21.2 |
| Exp. 4 | 2015_Jan8_hiNAHCO3_Daph | | |  | 6 |  | 27.888 |  | Mean | 20.4 |
|  | estimated from other experiments and lab notes. | | | | | | |  | StDevP | 0.3 |
|  | *chart in data file is blank (Temp Control failed and killed all organisms* | | | | | | | | |  |
|  | *and disabled Manta)* | | |  |  |  |  |  | MIN | 19.8 |
|  |  |  | Mean |  | 6.3 |  |  |  | MAX | 22.9 |
